# Tubules, rods and spirals: diverse modes of SepF-FtsZ assembling

**DOI:** 10.1101/2024.07.18.604210

**Authors:** Jagrity Choudhury, Barnali N. Chaudhuri

## Abstract

Z-ring formation by FtsZ, the master assembler of divisome, is a key step in bacterial cell division. Both formation and membrane anchoring of the Z-ring requires assistance of a number of Z-ring binding proteins, such as FtsA, EzrA, SepF, SepH and ZipA. SepF participates in bundling and membrane anchoring of FtsZ in gram-positive bacteria. We report *in vitro* biophysical studies of the interactions between FtsZ and cytoplasmic component of cognate SepF from three different bacteria: *Mycobacterium tuberculosis*, *Staphylococcus aureus* and *Enterococcus gallinarum.* While the cytosolic domain of SepF from *M. tuberculosis* is a dimer, those from *S. aureus* and *E. gallinarum* polymerize to form ring-like structures. Mycobacterial SepF helps in bundling of FtsZ filaments to form thick filaments and large spirals. On the other hand, ring-forming SepF from the *Firmicutes* bundle FtsZ into tubules.

## Introduction

Divisome, which is a gigantic, dynamic conglomerate of proteins spanning cytoplasm, membrane and periplasm, is the main coordinator of bacterial cell division (Cameron and Margolin, 2024, Haeusser and Margolin, 2016, Lutkenhaus et al., 2012, Naha et al., 2023). Divisome assembling is initiated at the cell division site by the formation of Z-ring from treadmilling FtsZ protofilaments, which requires assistance from several Z-ring associated proteins (Bisson-Filho et al., 2017, Huang et al., 2013, Squyres et al., 2021, Xiao and Goley, 2016, Yang et al., 2017). Z-ring recruits a number of early and late division proteins in a sequential manner for cell division septum formation (Cameron and Margolin, 2024, Gamba et al., 2009, Lutkenhaus et al., 2012). FtsZ contains an N-terminal globular region and a C-terminal flexible region (Heucas et al., 2020, Löwe and Amos, 1998, Matsui et al., 2012, Ruiz et al., 2022). The C-terminal part of FtsZ consists of an intrinsically disordered linker of variable length, followed by a conserved C-terminal tail region, and another short variable region (Buske and Levin, 2012, Heucus et al., 2017). The C-terminal tail serves as an interaction site for many FtsZ interacting proteins, including those that anchor FtsZ to the membrane (Cameron and Margolin, 2024, Cendrowicz et al., 2012, Haney et al., 2001, Mosyak et al., 2000, Radler and Loose, 2024, Singh et al., 2007). FtsZ polymerize in a “head-to-tail” fashion, and the GTPase active site of FtsZ is formed at the inter-subunit interface of the polymer (Wagstaff et al., 2017). How the polymers of FtsZ further bundle for Z-ring formation is not completely understood.

A number of proteins help in the bundling of FtsZ protofilaments to form the discontinuous-patchy Z-ring (Dajkovic et al., 2010, Durand-Heredia et al., 2011, Durand-Heredia et al., 2012, Gueiros-Filho and Losick, 2002, Hale and de Boer, 1997, Hale et al., 2011, Huang et al., 2013, Merino-Salomón et al., 2024, McQuillen and Xiao 2020, Ortiz et al., 2016, Rahman et al., 2020, Ramos-León et al., 2021, Rowlett and Margolin, 2015). Major FtsZ interacting proteins include FtsA (Schwedziak et al., 2012, Yan et al., 2000), ZipA (Haney et al., 2001, Mosyak et al., 2000), ZapA (Low et al., 2004), EzrA (Singh et al., 2007), SepF (Cendrowicz et al., 2012, Gundoğdu et al, 2011) and SepH (Ramos-Leon et al., 2021). Some of these FtsZ-bundling proteins are ubiquitous, and others are species-specific (Huang et al., 2013, Ortiz et al., 2016). Bundling of FtsZ filaments is critical for a properly functional Z-ring formation (Squyres et al., 2021). Unbundled, dispersed FtsZ filaments cannot recruit key divisome proteins, which in turn impairs septum formation (Squyres et al., 2021). Since FtsZ lacks a direct membrane-interacting domain, proteins like FtsA, ZipA and SepF serve as membrane anchors for the dynamic Z-ring (Barrows and Goley, 2021, Duman et al., 2013, Hale and Boer, 1997, Naha et al., 2023, Pichoff and Lutkenhaus, 2005)

SepF, which is a conserved protein in gram-positive bacteria, bundles and tethers FtsZ filaments to the membrane (Cendrowicz et al., 2012, Dey and Zhou 2022, Gola et al., 2015, Hamoen et al., 2006, Ishikawa et al., 2006, Naha et al., 2023, Singh et al., 2008). The absence of commonly occurring membrane anchors like FtsA and ZipA in mycobacteria makes SepF the sole membrane anchor of FtsZ and an essential cell division protein (Dey and Zhou 2022, Gola et al., 2015, Hamoen et al., 2006, Ishikawa et al., 2006). The soluble, C-terminal domain of SepF interacts with FtsZ in the cytoplasm, and an amphipathic helix at the N-terminus of SepF interacts with the membrane (Duman et al., 2013). The C-terminal domain of *Bacillus subtilis* SepF (bsSepF) polymerizes to form rings with ∼ 50 nm diameter, which can help bundle *B. subtilis* FtsZ (bsFtsZ) filaments into tubules (Duman et al., 2013, Gündoğdu et al., 2011). Binding to bsFtsZ leads to stacking of the bsSepF rings to aid tubule formation (Zhang et al., 2022). *In-vitro* bundling of FtsZ filaments by SepF is reported in mycobacteria but no tubular structures were reported (Bhattacharyya et al., 2015, Pende et al., 2021, Sogues at al., 2020). SepF from *Corynebacterium glutamicum* (cgSepF) bundles cognate FtsZ (cgFtsZ) into thick filaments instead of tubules (Sogues et al., 2020). Apart from FtsZ bundling and membrane anchoring, SepF is suggested to have a role in controlling the thickness of the nascent septum by forming arc-shaped polymers to cap the growing tip of the septum (Wenzel et al., 2021). Archaeal SepF preserved many structural features found in bacterial SepF, with distinct differences on FtsZ-SepF interaction (Pende et al., 2021). Recently, it was shown that SepF and FtsZ, both of which are members of Last Universal Common Ancestors, are capable of forming primitive Z-ring (Gulsoy et al., 2024).

Oligomeric forms of SepF, and SepF-FtsZ bundles, provided critical clues about the mechanisms of SepF-mediated FtsZ bundling and membrane tethering in *B. subtilis* (Gündoğdu et al., 2011) and *C. glutamicum* (Sogues et al., 2020). We present *in vitro* biophysical characterizations of SepF assembling and SepF-induced FtsZ bundling in three different bacterial species: *Mycobacterium tuberculosis*, *Staphylococcus aureus* and *Enterococcus gallinarum*. Our results reveal an interesting pattern, in which one group of ring-forming SepF tubulates FtsZ, while the other group of primarily dimeric SepF appears to laterally bundle FtsZ.

## Materials and Methods

### Cloning, expression and purification of proteins

The genes encoding the C-terminal domains of SepF from *S. aureus* (saSepF-CTD, residues 96-187) and *E. gallinarum* (egSepF-CTD, residues 115-204) were amplified by polymerase chain reaction (PCR) from the genomic DNA (gifted by Microbial Type Culture Collection and Gene Bank (MTCC), catalog numbers 1430^T^, 7049) and cloned in mcs1 of modified pET-Duet1 vector that contains TEV protease recognition site after the N-terminal hexa-histidine tag (pET-Duet1-TEV). Site directed mutagenesis was performed to generate point mutations of saSepF-CTD (saSepF-CTD_G145K_) and egSepF-CTD (egSepF-CTD_G163K_). The FtsZ gene from *S. aureus* (saFtsZ) and *E. gallinarum* (egFtsZ) was amplified in the same way and cloned into pET-Duet1-TEV and pET-28c respectively. Both the constructs of *M. tuberculosis*, FtsZ (tbFtsZ) and SepF-CTD (tbSepF-CTD, residues 146-241) used in this study, were commercially synthesized in codon-optimized form and cloned in pET-15b with N-terminal hexa-histidine tag (GenScript). These codon-optimized constructs were transformed into *Escherichia coli* BL21 (DE3) (Novagen) cells for recombinant protein expression. For the rest of the constructs, the vectors containing the gene of interest were transformed into chemical competent *E. coli* Rosetta (DE3) cells (Novagen). List of constructs, vectors, cloning sites and primers used in this work are listed in **supplementary tables S1-2**. The colonies obtained after transformation were used to inoculate starter cultures in the presence of appropriate antibiotics (100 μg/mL ampicillin for pET-15b/pET-Duet-TEV and 30 μg/mL kanamycin for pET-28c. Additional 35 μg/mL chloramphenicol was added during expression in Rosetta cells).

For large scale protein expression, 1 L of LB media was inoculated by 10 mL of primary culture and allowed to grow in the presence of antibiotics at 37 °C until OD_600_ reached 0.6. Next, the protein expression was induced with isopropyl-β-D-thiogalactoside (IPTG, 0.3 mM for SepF-CTD and 1 mM for FtsZ) and kept for overnight incubation at 16 °C with constant shaking. Harvested cells were stored at - 80 °C.

For purification of all the SepF-CTD constructs, the frozen cell pellets were thawed in buffer containing 25 mM Tris.HCl at pH 8.5, 100 mM NaCl, 1 mM ethylenediaminetetraacetic acid (EDTA) and 5% glycerol (buffer A). 10 μg/mL of Dnase I (Roche) was added to remove nucleic acid contamination. Cells were lysed by sonication followed by high-speed centrifugation at 13000 r.p.m. for 45 minutes. The supernatant was passed through IMAC sepharose beads charged with nickel and equilibrated with buffer A along with 10 mM imidazole. This is followed by a washing step. The protein was eluted by increasing the imidazole concentration in a step-wise manner. Next, fractions containing eluted proteins were concentrated, and injected to a pre-equilibrated size exclusion column (SEC, HiLoad 16/60 Superdex 200, GE Healthcare) equilibrated with buffer A. SDS-PAGE gels were used to evaluate protein purity at each step of purification. The pure proteins were concentrated and used for different experiments.

Purifications of all the FtsZ constructs used in this study were performed using the same protocol as mentioned above with different buffers. For saFtsZ and egFtsZ purification, buffer containing 50 mM Tris.HCl at pH 7.9, 100 mM NaCl, 1 mM EDTA and 10% glycerol (buffer B) was used. On the other hand, purification of tbFtsZ was performed in buffer containing 50 mM 4-(2-hydroxyethyl)piperazine-1-ethanesulfonic acid (HEPES) at pH 7.9, 200 mM KCl, 1 mM EDTA and 10% glycerol (buffer C).

### Negative stain Transmission Electron Microscopy

For SepF-CTD ring visualization, proteins were diluted to 100 μM immediately before grid preparation in a buffer containing 50 mM HEPES at pH 7.9, 200 mM KCl and 10 mM MgCl_2_. 300 mesh carbon coated copper grids (Agar Scientific) were glow discharged for 30 sec at 15 mA using the PELCO easiGlowTM 91000 Glow Discharge Cleaning System (Ted Pella, Inc.). 5 μL of protein was added to the glow-discharged grid, dried and stained with 2% uranyl acetate. The grids were imaged using JEM 2100 electron microscope (JEOL) at a voltage of 200 kV (in-house facility). To study the SepF induced FtsZ bundles, an equimolar ratio of FtsZ (12 μM) and SepF-CTD (12 μM) were mixed in polymerization buffer (mentioned below), incubated on ice for about 10 min, followed by addition of 1 mM of GTP and incubation for 10 min at 30 °C. Control experiments were done without adding SepF-CTD. The polymerization buffer used for saFtsZ and tbFtsZ bundling studies was buffer D (50 mM HEPES at pH 7.9, 200 mM KCl and 10 mM MgCl_2_). For egFtsZ bundling studies, buffer E (25 mM 2-(N-morpholino) ethanesulfonic acid (MES).KOH at pH 6.5, 50 mM KCl and 5 mM MgCl_2_) was used (Chen et al., 2007, Mukherjee and Lutkenhaus, 1999). All size measurements were performed using Fiji software (Schindelin et al., 2012).

### Analytical ultracentrifugation

Analytical ultracentrifugation (AUC) experiment was performed using in-house Beckman-Coulter XL-A analytical ultracentrifuge equipped with a 8 hole TiAn50 rotor. Two channel epon centerpieces were used with quartz windows. The experiments were performed at 40000 r.p.m. at 20 °C at three different protein concentrations. Buffer A was used for all the AUC experiments. The absorbance scans were measured at 280 nm. SEDNTERP (Hayes et al., 1995) was used for calculating the solvent density and solvent viscosity. Data fitting was done using the continuous distribution c(s) model by SEDFIT (Schuck et al., 2000).

### Small angle X-ray scattering

Small angle X-ray scattering (SAXS) experiment was performed at BM29 beamline, European Synchrotron Radiation Facility (ESRF), Grenoble, France. Purified tbSepf-CTD protein aliquots (3.6 mg/mL) along with background buffer (25 mM Tris.HCl at pH 8.5, 1 mM EDTA, 100 mM NaCl, and 10% glycerol) were frozen under liquid nitrogen and shipped in a dry shipper to the synchrotron site (Pernot et al., 2013). With a sample to detector distance was 2.9 m and X-ray wavelength of 1.0 Å, up to 10 acquisitions were collected with 0.5 s *per* frame. SAXS data was collected at three different concentrations. Data analysis was performed using the ATSAS 3.0 software suite (Manalastas-Cantos et al., 2021). Theoretical scattering profiles were calculated using CRYSOL, with default parameters (Franke et al., 2017). All graphs were prepared using Excel (Microsoft).

### Isothermal titration calorimetry

Interaction between tbSepF-CTD and a chemically synthesized 18-residue peptide from the C-terminal region of tbFtsZ (LSIGGDDDDVDVPPFMRR, GL Biochem) was examined using Isothermal titration calorimetry (ITC) in the in-house AUTOiTC 200 (Malvern) instrument. 87 μM of purified tbSepF-CTD was titrated against 1 mM of the C-terminal peptide with constant stirring at 750 r.p.m. 35 injections were given in total with injection volume of 0.8 μL (first injection 0.1 μL) and reference power of 10 μcal/s. A gap of 180 s was kept between two injections with filter period of 5 s. The experiment was performed in triplicates at 37 °C. The data was analyzed using the ORIGIN 6.0 software suite (Origin Lab).

### Structural modeling

The structures of saSepF-CTD, egSepF-CTD and tbSepF-CTD dimers were predicted using the Alphafold multimer option of ColabFold (Jumper et al., 2021, Mirdita et al., 2022). Helix packing angle at the A-interface was calculated in PyMOL (The PyMOL Molecular Graphics System, Schrodinger, LLC) using the plugin anglebetweenhelices.py (https://raw.githubusercontent.com/Pymol-Scripts/Pymol-script-repo/master/anglebetweenhelices.py). ConSurf (Chorin et al., 2020) and PyMol were used for preparation of figures. PyMOL was used for superposition of structures.

### Bioinformatic analysis

For performing the multiple sequence alignment (MSA), sequences of SepF homologs were selected from 18 different bacterial species and generated using the alignment tool Constraint based Multiple Alignment tool (COBALT) (Papadopoulos et al., 2007). MEGA version 11.0.11 (Tamura et al., 2021) was used for generating the phylogenetic tree using neighbor-joining method. Jalview (Waterhouse et al., 2009) was used to generate the MSA shown in the figures.

### Data availability statement

SAXS data are deposited in Small Angle Scattering Biological Data Bank (SASBDB, www.sasbdb.org; accession codes: SASDUX2, SASDUY2 and SASDUZ2). Remaining data are deposited to our institutional data repository, and can be obtained from the corresponding author upon request.

## Results and discussion

### 1. Cytosolic components of saSepF and egSepF form ring-like structures with conserved interfaces

To characterize the oligomeric forms of saSepF-CTD and egSepF-CTD, negative staining electron microscopy was performed. Both saSepF-CTD and egSepF-CTD appears to form large rings (**Figure 1A-D**). The inner diameters of these rings were ∼ 35 nm and ∼ 32 nm for saSepF-CTD and egSepF-CTD, respectively (**Figure 1E, F; Table 1**). Widths of these rings were of the order of a few nm (**Figure E, F**).

**Figure 1.**
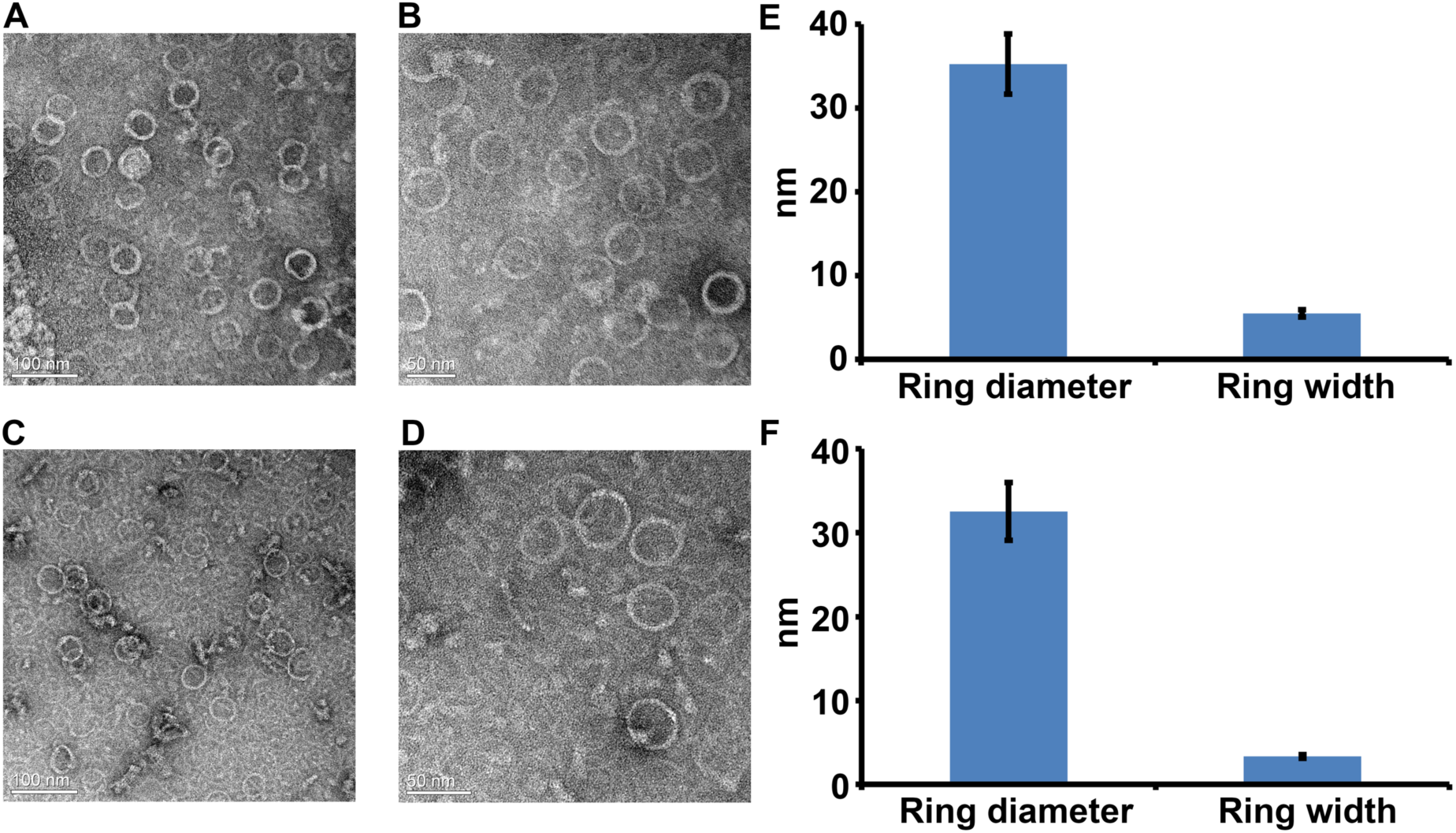
Ring-shaped polymers formed by saSepF-CTD and egSepF-CTD. Negative stain electron micrographs of (A-B) saSepF-CTD rings and (C-D) egSepF-CTD rings. Scale bars 50 or 100 nm. (E) Bar diagram showing measured inner diameter (N = 66) and width (N = 69) of saSepF-CTD rings. (E) Bar diagram showing measured and inner diameter (N = 112) and width (N = 69) of egSepF-CTD rings.

**Table 1.** Sizes of the SepF and SepF-FtsZ polymers. All measurements were obtained from the electron micrographs using the Fiji software (Schindelin et al., 2012).

| Species | Average (inner) diameter of the SepF-CTD ring | Average (inner) diameter of FtsZ tubules (nm) | Average width of FtsZ tubules (nm) | Average (outer) diameter of the SepF-CTD ring |
| --- | --- | --- | --- | --- |
| *S. aureus* | 35.2 ± 3.6<br>(N = 66) | 37.7 ± 3.6<br>(N = 38) | 6.4 ± 1.7<br>(N = 38) | 43.5 ± 3.0<br>(N = 42) |
| *E. gallinarum* | 32.6 ± 3.4<br>(N = 112) | 36.0 ± 7.5<br>(N = 27) | 8.5 ± 4.0<br>(N = 27) | 37.8 ± 3.6<br>(N = 62) |

Two inter-subunit interfaces, one formed by two α-helices (the A-interface) and another by β-strands (the B-interface), were described as two distinct polymerization interfaces for ring formation in the cytosolic domain of SepF from *B. subtilis* (bsSepF-CTD, **Figure 2A**) (Duman et al., 2013). A conserved glycine residue at the A-interface (G109, *B. subtilis* numbering; Duman et al., 2013) was previously reported to be critical for polymerization of bsSepF-CTD, as it allows close packing of the α-helices at the intermolecular interface (**Figure 2A, B**). Alphafold2 predicted structures of dimeric saSepF-CTD and egSepF-CTD were similar to the structure of bsSepF-CTD (**Figure 2C, D, E**). The B-interface is the dimerization interface in the Alphafold2 predicted dimeric structures of saSepF-CTD and egSepF-CTD (**Figure 2C, D**). The A-interfaces in these predicted structures of saSepF-CTD and egSepF-CTD are available for further polymerization. To find out whether saSepF-CTD and egSepF-CTD polymerize *via* this A-interface, we mutated the conserved glycine residues with lysine (saSepF-CTD_G145K_ and egSepF-CTD_G163K_, **Figure 2C, D**). Intermolecular packing of the alpha helices at the A-interface will be obstructed by the bulky side chain of lysine, which is expected to impair ring formation. Since negative stain electron micrographs obtained from none of these two mutants revealed ring formation, we performed AUC experiments. AUC data of both the mutant proteins show major population of dimer, as expected (**Figure 2F, G**). For saSepF-CTD_G145K_, a prominent peak was seen at s_20, w_ of 2.4±0.19S with a frictional ratio of 1.2±0.05. For egSepF-CTD_G163K_ mutant, a major peak was seen at s_20, w_ of 1.6±0.09S with a frictional ratio of 1.8±0.04. The molecular weights were estimated to be 23.8 kDa for saSepF-CTD_G145K_ and 23.1 kDa for egSepF-CTD_G163K_, which are close to the corresponding dimeric weights of the proteins (26.2 kDa for saSepF-CTD_G145K_ and 25.4 kDa for egSepF-CTD_G163K_ respectively). These dimers are understandably formed *via* the B-interface of SepF-CTD, and they cannot further oligomerize for ring formation due to the A-interface-obstructing mutations. Our mutational study supports that the A-interface is involved in ring formation of saSepF-CTD and egSepF-CTD.

**Figure 2.**
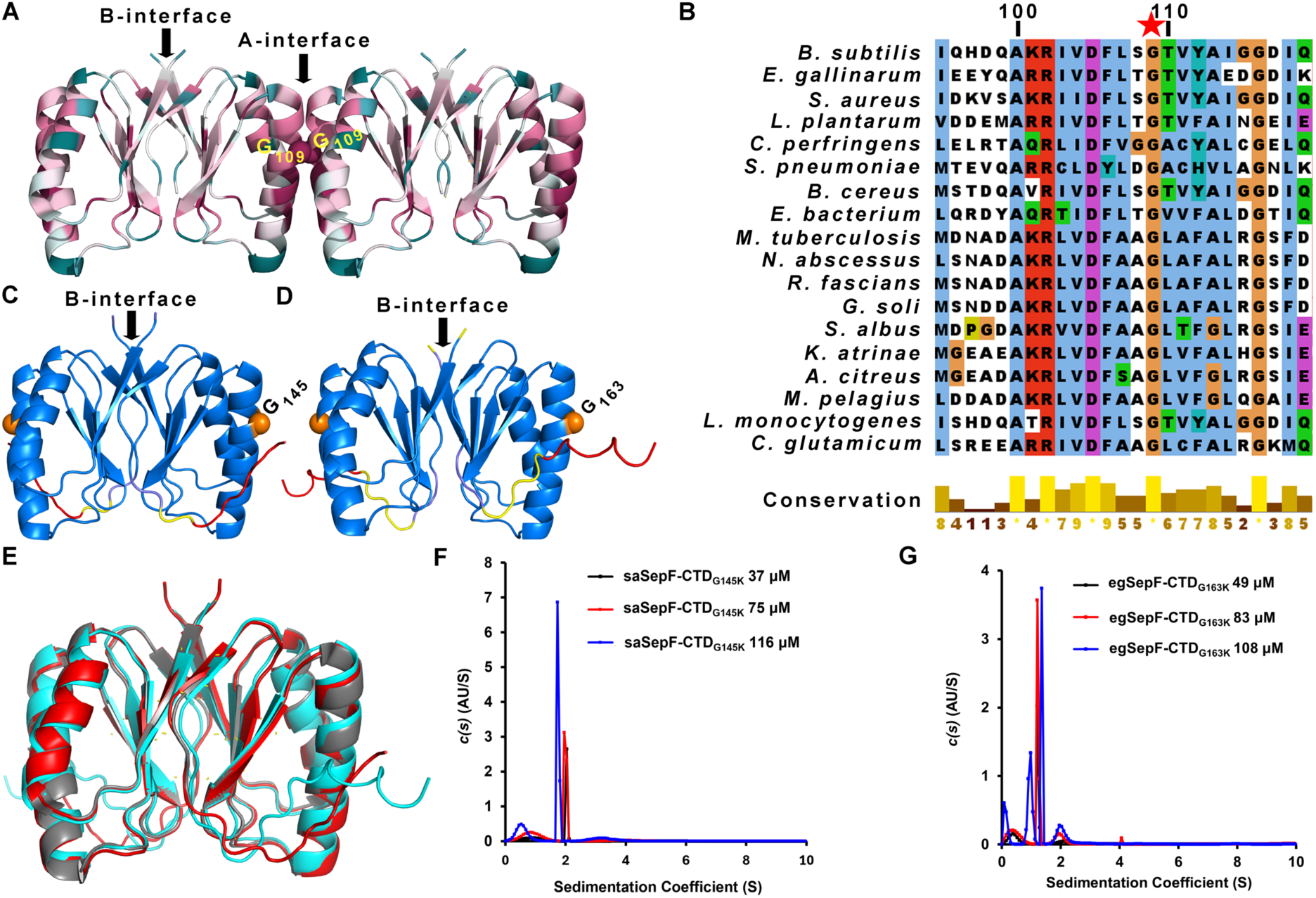
Polymerization interfaces for ring formation are conserved in bsSepF, saSepF and egSepF. (A) Cartoon of tetrameric bsSepF-CTD (PDB ID 3ZIH), coloured according to sequence conservation score (Consurf, Chorin et al., 2020), is shown. The two proposed polymerization interfaces, the A-interface and the B-interface, are marked. The conserved glycine residue (G109) at the A-interface is labelled and shown in CPK. (B) Multiple sequence alignment of SepF sequences from different bacterial species, showing the region encompassing the A-interface (prepared using Jalview, Waterhouse et al., 2009). The conserved glycine residue from the A-interface is marked with a star. Predicted models of (C) saSepF-CTD and (D) egSepF-CTD dimers, coloured according to the pLDDT scores (blue; pLDDT> 90 to red; pLDDT< 50). The mutated glycine residue is shown in orange CPK. (E) Superposed predicted models of saSepF-CTD (red), egSepF-CTD (cyan) and crystal structure of bsSepF-CTD (grey, PDB ID 3ZIH) are shown as cartoons. The RMSD between bsSepF-CTD and saSepF-CTD is 0.4 Å (143 Cα atoms), and between bsSepF-CTD and egSepF-CTD is 0.8 Å (157 Cα atoms). (F-G) AUC data showing sedimentation co-efficient distribution profiles of (F) saSepF-CTD_G145K_ and (G) egSepF-CTD_G163K_ at three different concentrations.

Our efforts to purify B-interface obstructing mutants were not successful and hence the role of the B-interface in ring formation could not be experimentally verified.

### 2. tbSepF-CTD is primarily a dimer in solution

Although tbSepF-CTD is reported to form rings (Sogues et al., 2020, Wenzel et al., 2021), we were unable to locate abundant rings for tbSepF-CTD in our TEM experiments. Only one ring was located in a TEM grid (**Figure S1**), which could be an artefact. To determine the oligomeric state(s) of tbSepF-CTD in solution, we performed AUC and SAXS experiments at a range of concentrations.

The AUC profiles show that a major population of tbSepF-CTD exists as a dimer in solution, with an s_20, w_ of 2.12±0.04 at a frictional ratio of 1.53±0.02 (**Figure 3A**). The AUC-derived molecular mass of 27 kDa is comparable to the theoretical dimeric mass of tbSepF-CTD (24.8 kDa). The dimensionless Kratky plot obtained from SAXS data suggested that tbSepF-CTD is globular and folded (**Figure 3B**). The maximum particle diameter (D_max_) of the scattering particle obtained from the pair distribution function plot was ∼ 9 nm (**Figure 3C**). The Guinier analysis provided the average radius of gyration (R_g_) of ∼ 2.4 nm (**Figure 3D**). The estimated molecular mass of the scattering particle was ∼ 23.7 kDa (Bayesian inference; Hajizadeh et al., 2018), which corresponds to the dimeric state of tbSepF-CTD. All mass prediction tools available in the ATSAS software predicted mass values between ∼ 24 kDa – 30 kDa. Thus, both AUC and SAXS data were consistent with a dimeric form of tbSepF-CTD in solution.

**Figure 3.**
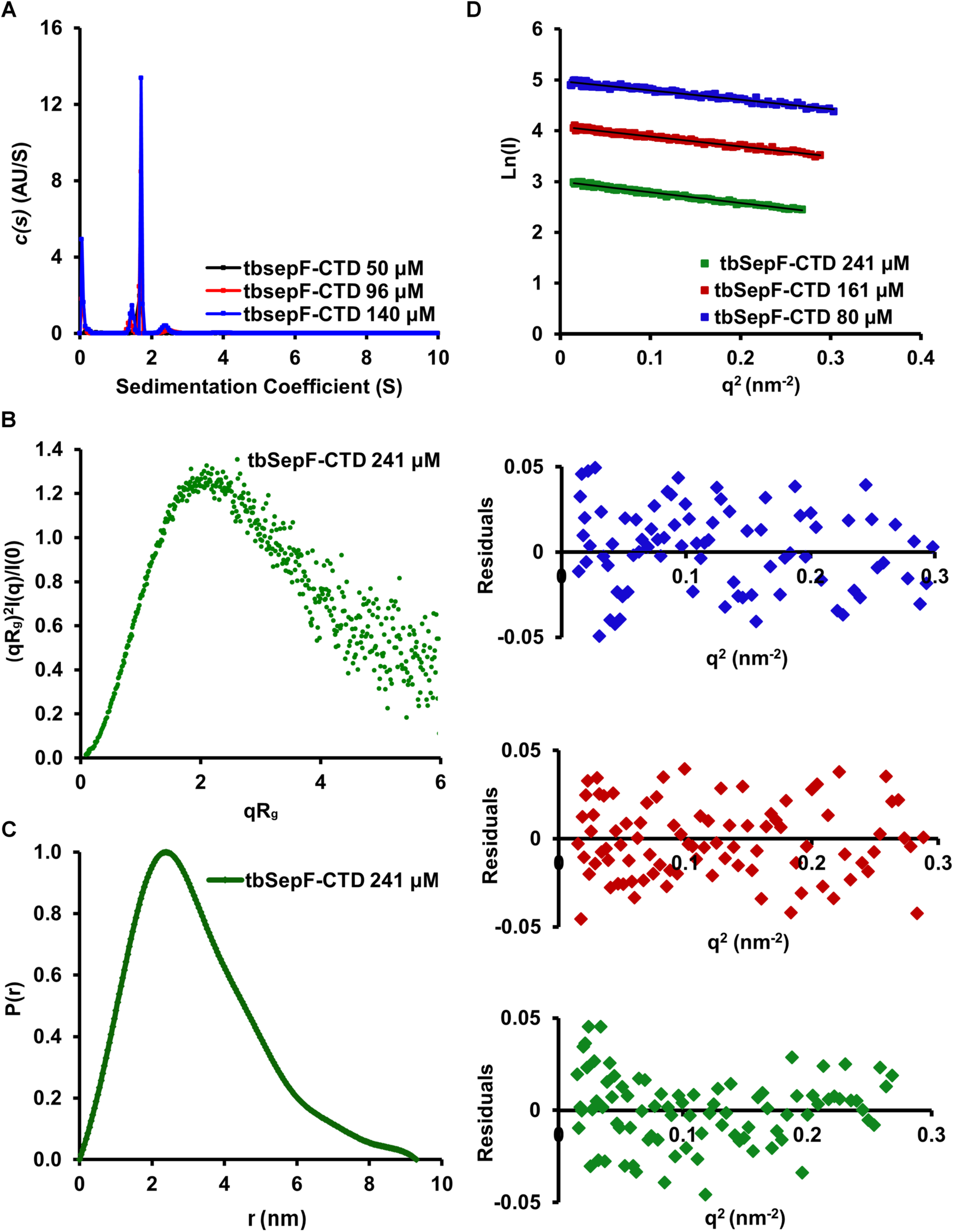
AUC and SAXS data supports that tbSepF-CTD is a dimer. (A) AUC data showing sedimentation coefficient distribution profile of tbSepF-CTD at three different concentrations. (B) Dimensionless Kratky plot (plot of (qR_g_)^2^(I(q)/I(0) *versus* qR_g_, where q is momentum transfer at nm^-1^, I(q) is the intensity in arbitrary unit, I(0) is forward scattering intensity, R_g_ is radius of gyration) and (C) pair-distribution function plot (P(r) *versus* r, where r is pairwise distance between atoms) obtained from the SAXS data of tbSepF-CTD at 241 μM concentration. The values on the Y-axis are normalized to maximum value of 1. (D) Guinier plot of tbSepF-CTD SAXS data at three different concentrations. Corresponding residual plots are shown in the lower panels. Plots are shifted along the y-axis for better visualization.

Next, we predicted the structure of tbSepF-CTD dimer using Alphafold multimer option in Colabfold (Jumper et al., 2021, Mirdita et al., 2022). The predicted, top-ranking dimeric model of tbSepF-CTD resemble the crystal structure of the C-terminal segment of SepF from *C. glutamicum* (cgSepF-CTD, PDB ID 6SCP, Sogues et al., 2020) with A-interface as the dimerization interface and an extra “α3 helix” at the C-terminus (**Figure 4A-B**). Inter-subunit packing angle between the two alpha helices (residues 163-180 in tbSepF) at the A-interface of this tbSepF-CTD model is comparable to the corresponding packing angle, which is ∼ 50°, in cgSepF-CTD dimer (**Figure 4B**). Although both A-interface and B-interface regions appear to be conserved in actinobacterial SepF (**Figure 4C**), none of the predicted structures of tbSepF-CTD dimerized with the B-interface. A theoretical scattering curve generated from the top-ranked predicted model of dimeric tbSepF-CTD, which was formed using the A-interface, agreed reasonably with the experimental SAXS scattering curve, with a reduced χ^2^ value of 2.0 (**Figure 4D**). A B-interface dimer of tbSepF-CTD, which was prepared by superposing tbSepF-CTD subunits on the bsSepF-CTD dimer structure, does not agree well with the SAXS data (reduced χ^2^ value of 4.7, **Figure 4D**). Agreement between experimental SAXS data and predicted model of tbSepF-CTD further supports that tbSepF-CTD dimerize using the A-interface in the same way as cgSepF, with conserved packing geometry.

**Figure 4.**
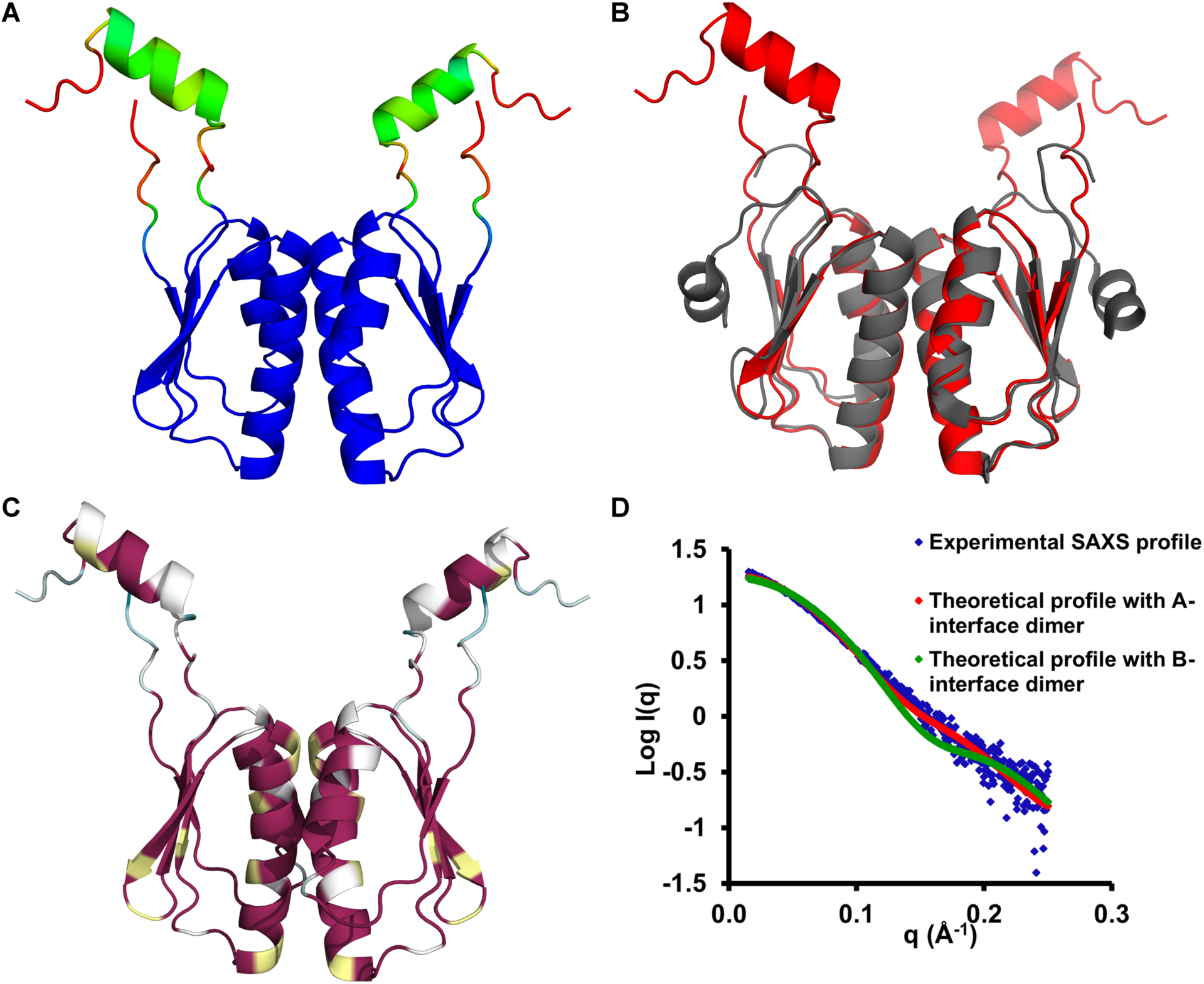
Predicted model of tbSepF-CTD dimer concurs with SAXS data. Cartoon diagrams of Alphafold2 predicted tbSepf-CTD model (A) coloured according to pLDDT score (blue; pLDDT> 90 to red; pLDDT< 50), (B) coloured in red, shown along with the superposed cgSepF-CTD structure (grey; PDB ID 6SCP; RMSD 0.54 Å for 65 Cα atoms), and (C) coloured according to sequence conservation in *Actinobacteria* (obtained using 1000 actinobacterial SepF sequences by ConSurf; Chorin et al., 2020). (D) Plot of logI *versus* q (I(q) is intensity in arbitrary unit and q is momentum transfer in Å^-1^) showing the fit between experimental SAXS data (blue diamond) and theoretical scattering profiles obtained from the predicted, A-interface dimer (red diamond) and B-interface dimer (green diamond) of tbSepF-CTD.

### 3. Ring-forming saSepF-CTD and egSepf-CTD tubulates cognate FtsZ

The effect of ring-forming SepF-CTD on the bundling of FtsZ filaments was studied using negative stain TEM. The electron micrographs of saFtsZ in the presence of saSepF-CTD and GTP showed the formation of long tubular structures (**Figure 5A-B**, **Table 1**). These tubules are similar to the previously reported tubules formed by bsFtsZ in the presence of cognate SepF and nucleotide (Gundoğdu et al., 2011). No tubules were observed for the control sample of saFtsZ in the absence saSepF-CTD (**Figure 5C**). At times, we observed rings with diameters similar to that of saSepF-CTD rings in the background (**Figure 5A**). Similar to saFtsZ, egFtsZ formed tubules in the presence of egSepF-CTD (**Figure 5D-E**, **Table 1**) that was absent in the egFtsZ control (**Figure 5F**). The inner diameters of these saFtsZ and egFtsZ tubules (**Figure 5G, H**) are approximately similar to the outer diameters of the cognate SepF-CTD rings (**Figure 1E-F**; **Table 1**). SepF, however, does not form a full circular ring structure inside the cell, and forms a semi-circular polymeric structure to cap the septum (Duman et al., 2013). Earlier, it was proposed that bsSepF rings stack inside the tubules, and bsFtsZ filaments are aligned on the outer part of the tubules (Gundoğdu et al., 2011). Stacking of SepF rings upon addition of the C-terminal peptide segment of FtsZ has been reported (Zhang et al., 2022). Probably the FtsZ-interacting saSepF-CTD and egSepF-CTD rings stack and align the FtsZ filaments on the outer rim of the stacked rings for tubule formation in the same manner.

**Figure 5.**
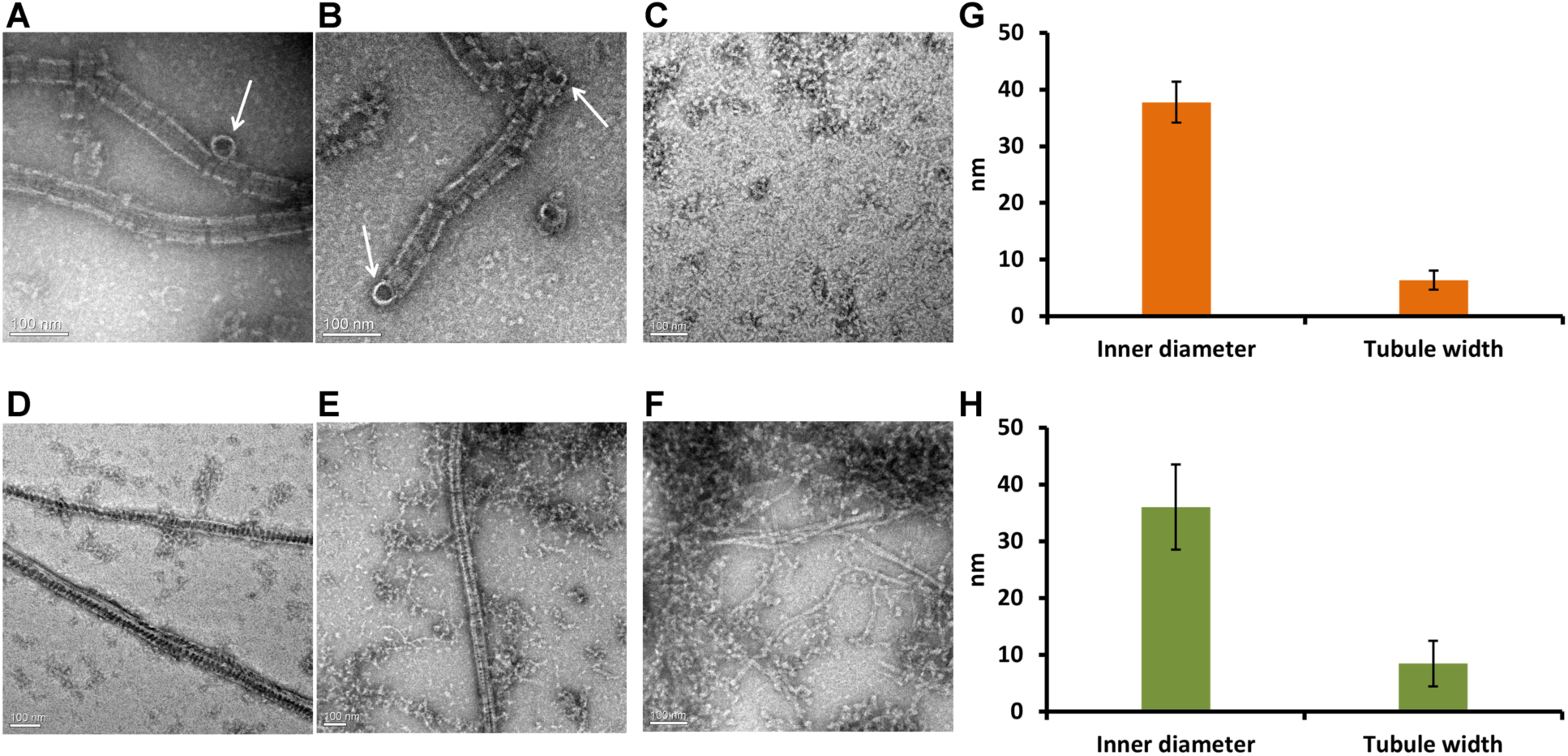
saSepF-CTD and egSepf-CTD tubulates cognate FtsZ. Negative stain electron micrographs of (A-B) saFtsZ tubules in the presence of saSepF-CTD and GTP, and (C) saFtsZ in the presence of GTP. saSepF-CTD rings located next to the tubules are marked with white arrows. Negative stain electron micrographs of (D-E) egFtsZ tubules in presence of egSepF-CTD and GTP, and (F) egFtsZ in presence of GTP. (G-H) Bar diagram showing (G) the measured saFtsZ inner tubule diameter (N = 38) and width (N = 39), and (H) the measured egFtsZ tubule diameter (N = 27) and width (N = 27). Scale bar 100 nm.

### 4. tbSepF bundles tbFtsZ into rod-shaped thick filaments and spirals

To find the effect of dimeric tbSepF-CTD on tbFtsZ bundling, we performed negative stain TEM. Electron micrographs showed thick filaments, with width of about 19 ± 5.6 nm, formed by tbFtsZ in the presence of tbSepF-CTD and GTP (**Figure 6A-B**). Similar thick filaments were reported for cgFtsZ bundled by its cognate SepF (Sogues et al., 2020). On the other hand, tbFtsZ in the absence of tbSepF-CTD formed thinner filaments (5.7 ± 2.0 nm), which is consistent with earlier reports (**Figure 6C**; Chen et al., 2007).

**Figure 6.**
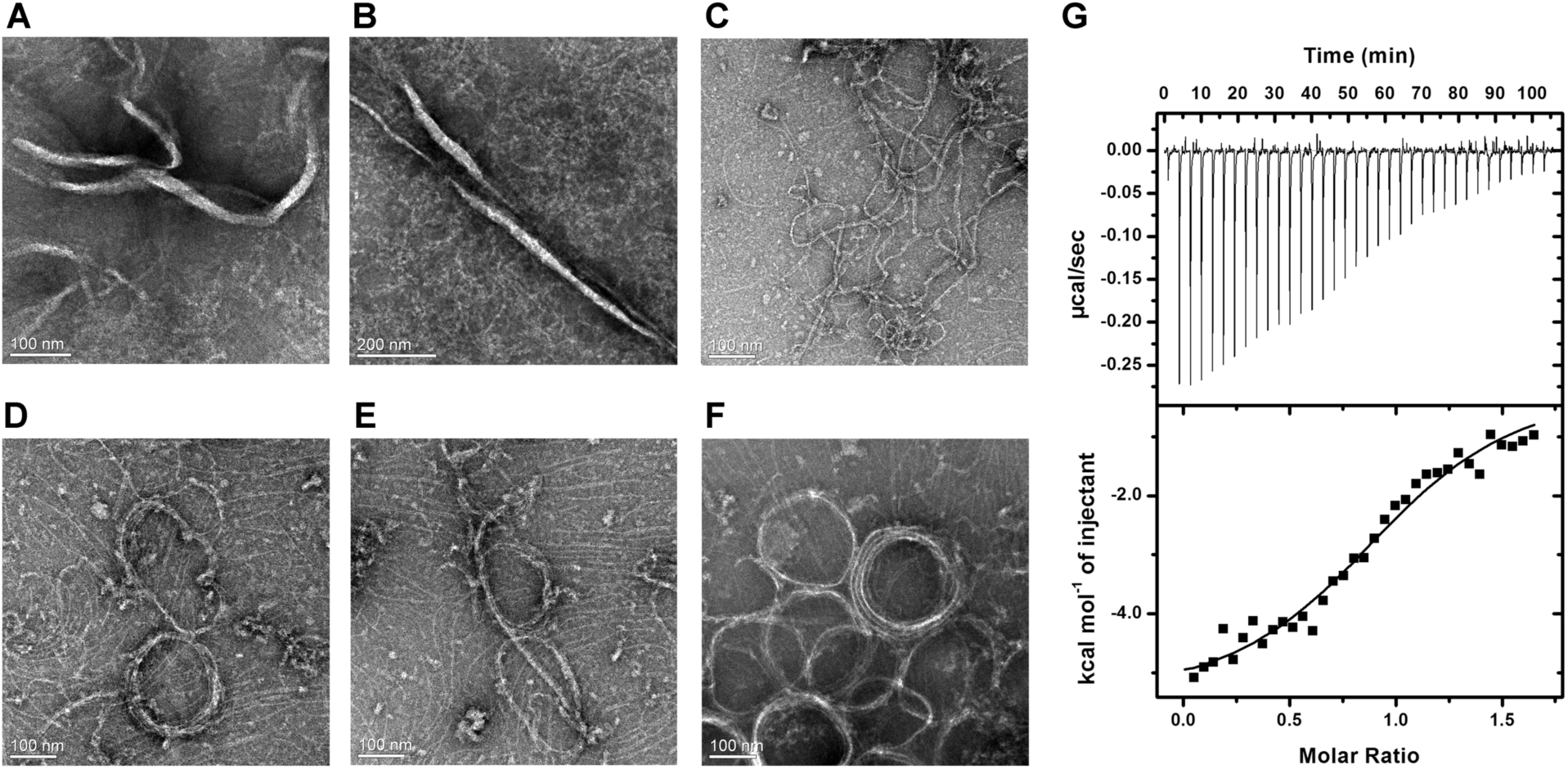
tbSepF-CTD bundles tbFtsZ into thick filaments and spirals. Negative stain electron micrographs showing (A-B) tbFtsZ thick filaments in the presence of tbSepF-CTD and GTP. Scale bar 100 or 200 nm. (C) Thin filaments of tbFtsZ in the presence of GTP. (D-F) tbFtsZ spirals in the presence of tbSepF-CTD and GTP. Scale bar 100 nm. (E) ITC thermogram (upper panel) and fit data (lower panel) showing interaction between tbSepF-CTD and an 18-meric peptide from the C-terminus of tbFtsZ.

Apart from the thick filaments, tbFtsZ formed large spirals in the presence of tbSepF-CTD and GTP (**Figure 6D-F**). The average diameter of these large ring-shaped structures was about 185.2 ±22 nm. For a few cases, longest dimensions of these spirals were > 250 nm. Similar spiral structures were reported for tbFtsZ in the presence of crowding agents with metal ions such as methyl cellulose with Rb^+3^ or K^+^ (Popp et al., 2009, Popp et al., 2010). Metal ions can possibly support this spiral formation by helping in the lateral association of bent filaments (Popp et al., 2010). Although our buffer contained K^+^ ions, tbFtsZ on its own did not form spirals in the absence of tbSepF-CTD. We considered the possibility that tbSepF-CTD act like a crowding agent to assist spiral formation in the presence of K^+^ ions and do not directly interact with tbFtsZ filaments. Therefore, we confirmed direct physical interaction between tbSepF-CTD and tbFtsZ (**Figure 6E**). The C-terminal region of tbFtsZ binds tbSepF-CTD with 1:1 stoichiometry, and dissociation constant (K_d_) of 19.7 μM (±7.3 μM). Similar dissociation constants were reported for interaction between FtsZ and other FtsZ-associated proteins such as FtsA (45-58 μM, Szwedziak et al., 2012), ZipA (35 μM, Mosyak et al., 2000), ZapD (10 μM, Choi et al., 2016) and SepF (15 μM, Sogues et al., 2020). In addition, crystal structure of cgSepF-CTD bound to the C-terminal peptide of cgFtsZ has been reported (Sogues et al., 2020). Since a dimeric tbSepF binds two molecules of tbFtsZ, the thick filaments and spirals are probably formed by lateral bridging of tbFtsZ filaments by tbSepF-CTD. It is well-known that GTP hydrolysis bends FtsZ filaments (Lu and Erickson, 1999). These large spirals are likely result of bending of the bundled straight tbFtsZ filaments owing to GTP hydrolysis.

Since it was proposed that the presence of the protruding helical region might block B-interface formation in cgSepF-CTD (Sogues et al., 2020), we prepared a construct of tbSepF in which this helical segment was removed (deleted residues 228-241, Δα-tbSepF-CTD). Purified Δα-tbSepF-CTD eluted at the same place as wild-type tbSepF-CTD in the size-exclusion column run, and induced tbFtsZ to form spiral structures in the presence of GTP (data not shown). However, further polymerization of membrane-anchored, full-length tbSepF dimers remains a possibility (Sogues et al., 2020).

## Conclusion

SepF is a bacterial cell division protein that interacts with FtsZ and performs multiple functions for smooth and accurate division of the cell. Our biophysical study of SepF and SepF-FtsZ assembling in three different bacteria (this work), combined with earlier reports (Duman et al., 2013, Gundoğdu et al., 2011, Sogues et al., 2020), reveal a pattern in which the oligomeric states of SepF are linked to how they bundle cognate FtsZ (**Figure 7**). SepF-CTD from three *Firmicutes*, *B. subtilis* (Duman et al., 2013), *S. aureus* and *E. gallinarum,* oligomerize to form rings (**Figure 7A**). All these ring-forming SepF-CTD can tubulate FtsZ, probably by arranging the FtsZ filaments around the outer circumference of the ring (Gundoğdu et al., 2011; this work). This ring forming property of SepF, which requires formation of both A-interface and B-interface, is conserved in a number of *Firmicutes* (Wenzel et al., 2021). On the other hand, actinobacterial SepF (*M. tuberculosis* and *C. glutamicum*) from a different branch of the phylogenetic tree do not form rings (this work; Sogues et al., 2020). tbSepF-CTD mostly form dimers and bundle FtsZ into thick filaments (**Figure 7**). In addition, we observed large spirals of tbFtsZ in the presence of tbSepF-CTD (**Figure 6D-F**; **Figure 7**), at times with sizes approaching the diameter of rod-shaped mycobacteria (300 – 500 nm, Cook et al., 2009). Similar shape formation by FtsZ in the presence of ZapD has been reported (Merino-Salomon et al., 2024). Although the biological significance of these spirals is not clear at present, these large structures might be relevant for primitive Z-ring formation. In conclusion, our study provided key insights into the diversity in the FtsZ-SepF bundling in different bacteria.

**Figure 7.**
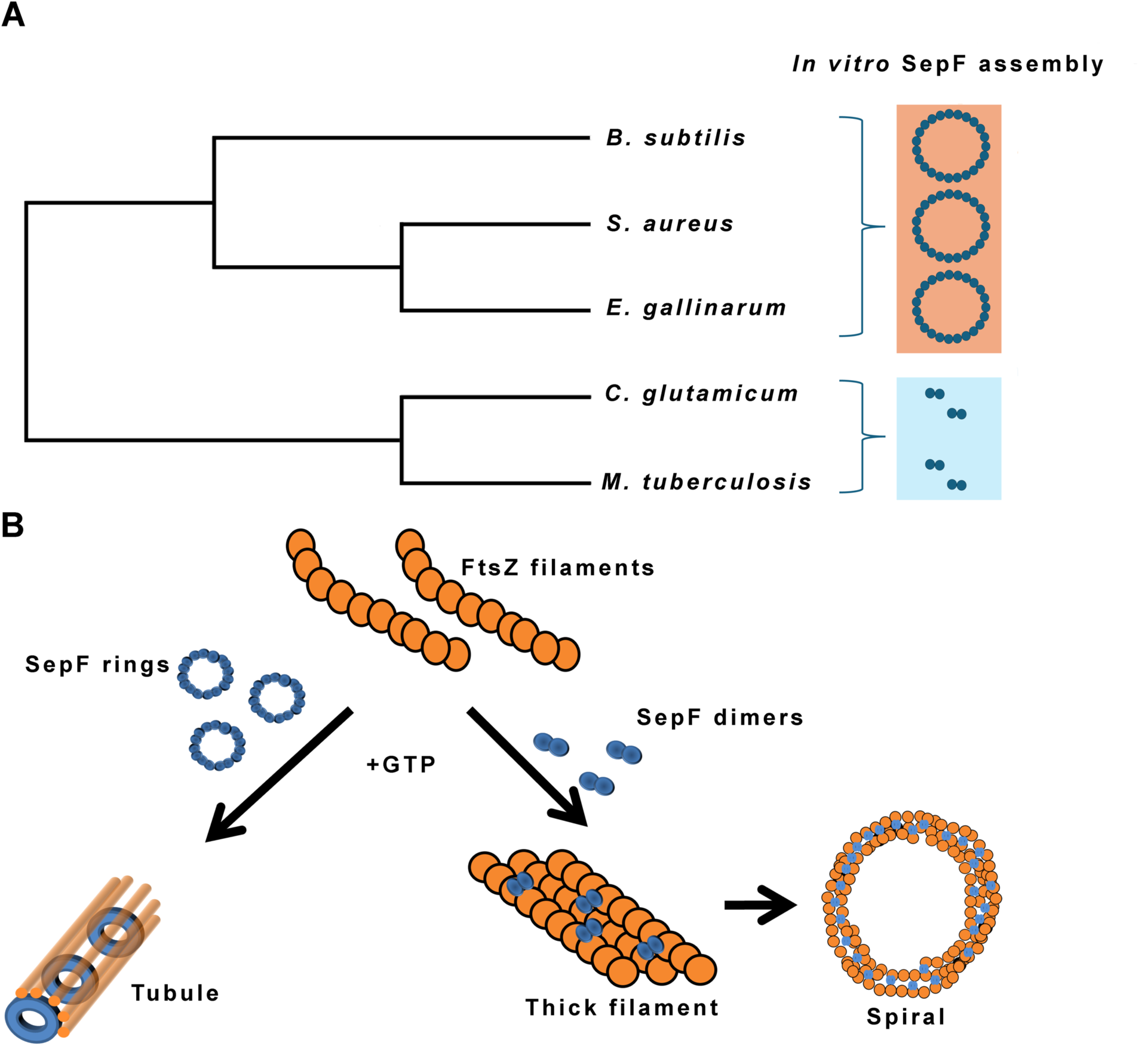
Diversity of SepF and SepF-FtsZ assembling. (A) A phylogenetic tree showing the evolutionary relations of SepF discussed in this study, along with schematics depicting their oligomeric states. (B) Schematic diagrams showing various modes of SepF-FtsZ assembling.

## Acknowledgements

The authors acknowledge CSIR for intramural funding and facilities. JC is grateful to the Department of Biotechnology (DBT), Govt. of India, for fellowship. The authors thank MTCC for providing the genomic DNA, and ESRF-access program supported by DBT and conducted by Regional Centre for Biotechnology for providing the beamtime for SAXS experiments. We thank the staff at beamline BM29, ESRF, for help with SAXS data collection.

## Supplementary material

**Table S1.** Constructs used in the study.

| Organism | Construct | Vector | Cloning site |
| --- | --- | --- | --- |
| *M. tuberculosis*<br>_H37Rv | N_term_6His_tev site_<br>tbSepF_CTD(146-241) | pET-15b | NcoI-XhoI |
| | N_term_6His_tev site_ $\Delta\alpha$-<br>tbSepF-CTD (146-227) | pET-15b | NcoI/XhoI |
|  | N_term_6His_tev site_<br>tbFtsZ | pET-15b | NcoI-XhoI |
| *S. aureus* | N-term_6His_tev<br>site_saSepF-CTD | pET-Duet1-TEV | NheI_HindIII |
|  | N-term_6His_tev site<br>_saSepF-CTD<sub>G145K</sub> | pET-Duet1-TEV | NheI_HindIII |
|  | N-term_6His_tev site<br>_saFtsZ | pET-Duet1-TEV | NheI_XhoI |
| *E. gallinarum* | N-term_6His_tev site<br>_egSepF-CTD | pET-Duet1-TEV | NheI_HindIII |
|  | N-term_6His_tev site<br>_egSepF-CTD<sub>G163K</sub> | pET-Duet1-TEV | NheI_HindIII |
|  | N-term_6His_tev site<br>_egFtsZ | pET-28c | NdeI_BamHI |

**Table S2.**
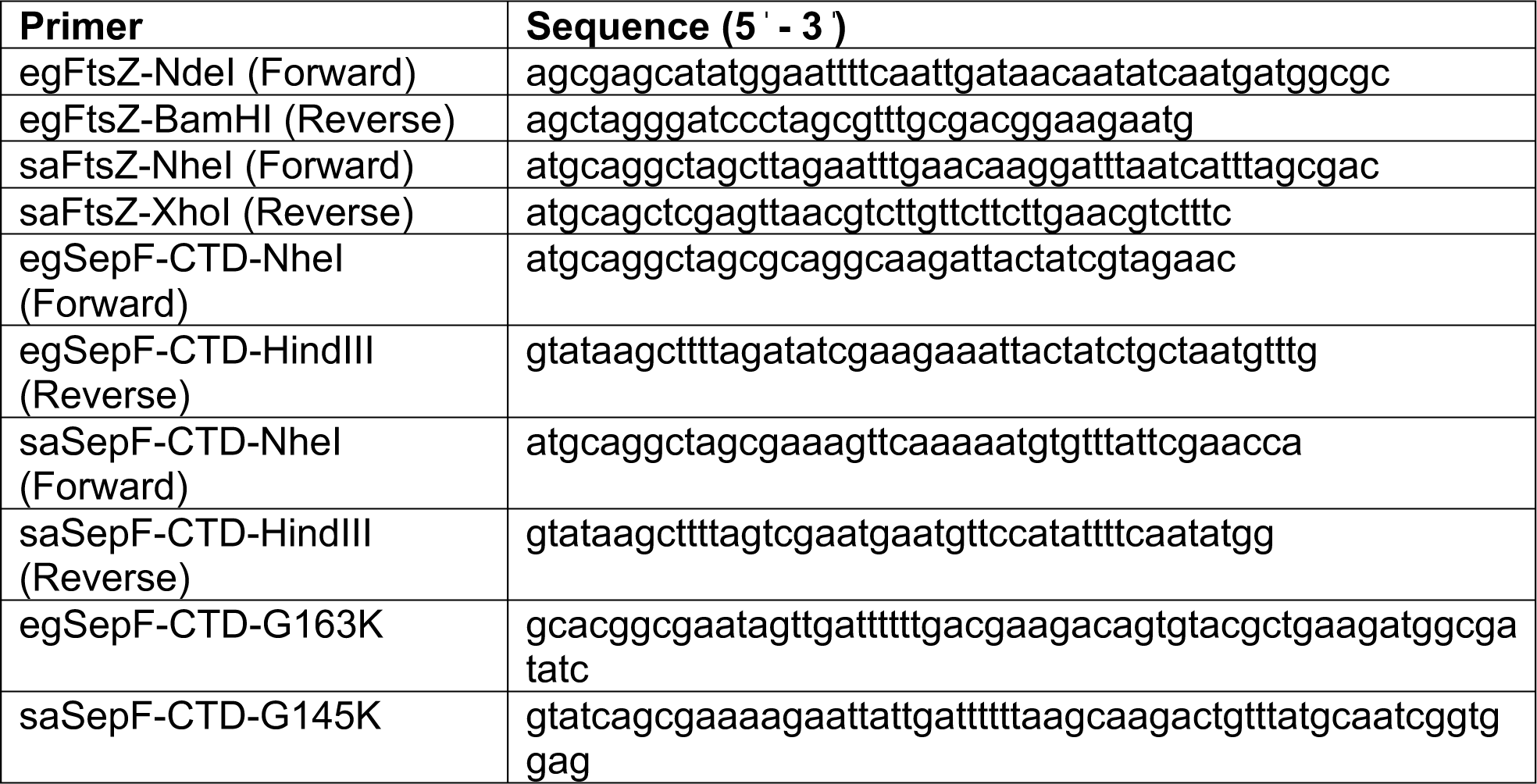
List of primers used in this study.

**Figure S1.**
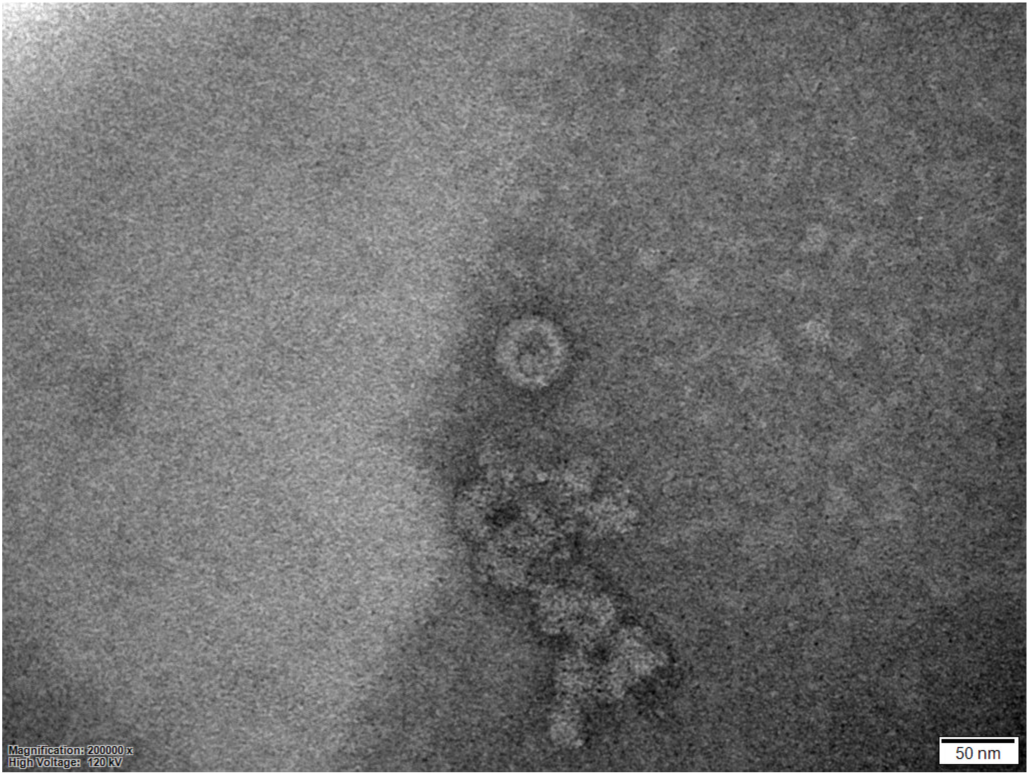
Negative stain electron micrograph of tbSepF-CTD. Scale bar 50 nm.

